# Programmable sequence-resolved *in situ* profiling of glycoRNA-protein interactomes by GlycoRNA-Map

**DOI:** 10.64898/2026.09.29.755375

**Authors:** Xinming Zhang, Yingying Zhu, Katelyn M. Baumer, Yao Lu, Yu Fu, Jiahao Liao, Xiangli Shao, Yuduo Chang, Eva Qiao, Haoyue Dong, Álvaro Sierra Valverde, Hsin-Chih Yeh, Shu-Hsia Chen, Hans Renata, Yuan Ma

## Abstract

Cell-surface glycoRNAs participate in diverse biological processes, but they were investigated as pooled populations and the protein interactome of individual glycoRNA species remain unresolved. Here we develop GlycoRNA-Map, a programmable photocatalytic proximity-labeling method that maps proteins surrounding sequence-defined endogenous glycoRNAs on living cells. A sialic-acid-binding aptamer and an RNA-hybridization probe jointly recruit a fluorophore-modified oligonucleotide through dual recognition of the glycan and RNA moieties of glycoRNAs. Light-activated biotinylation followed by quantitative mass spectrometry identified 200-418 enriched proteins for each of five glycoRNAs in A549 cells. Comparison of these proteomes revealed limited overlap among different glycoRNA sequences and across cell types, demonstrating that glycoRNA-associated protein environments depend on both RNA identity and cellular context. Transforming growth factor-β receptors 1 and 2 (TGFBR1 and TGFBR2) were identified in the proximal proteomes of multiple glycoRNAs. Removal of cell-surface RNAs enhanced TGF-β receptor signaling and promoted cell migration, demonstrating that glycoRNAs can functionally regulate extracellular signaling by interacting with their protein neighborhoods. GlycoRNA-Map thus provides an accessible strategy for resolving sequence-specific glycoRNA-protein interactome and discovering their roles in membrane signaling.

## Introduction

RNA has traditionally been viewed primarily as an intracellular molecule that regulates gene expression and protein synthesis^1, 2^. Recently, diverse RNA species have also been detected on the outer surfaces of living cells^3, 4^. Notably, some cell-surface RNAs are covalently modified with glycans and are referred to as glycoRNAs^3^. These molecules are found across multiple tissues, species, and cell types^3–22^. Functionally, glycoRNAs regulate immune-cell recruitment and cell-cell interactions^4, 5, 7, 23^, influence immune recognition and immune evasion^8, 24^, and facilitate the cellular entry of cell-penetrating peptides^10, 25^. GlycoRNAs have also been implicated in heparan sulfate-dependent signaling^11^, suppression of nuclear factor-κB signaling and inflammatory cytokine production in neurons^26^, and regulation of alveolar epithelial function^27^. Moreover, the abundance and composition of glycoRNAs can be utilized to distinguish cancer cells from nonmalignant cells and differentiate specific cancer types, suggesting their potential applications as disease-associated molecular signatures^5, 12, 13, 28^. Collectively, these discoveries suggest that glycoRNAs constitute a previously underappreciated regulatory layer at the plasma membrane.

RNAs need to interact with proteins to fully exert their functions, and proteins also bind to RNAs to act as regulators. It has now become clear that RNA-protein interactions play important roles in many biological processes underlying health and disease^29–35^. At the cell surface, glycoRNAs are thought to exert their functions through interactions with proximal proteins. For example, glycoRNAs form nanoclusters with cell surface RNA-binding proteins, including cell-surface nucleophosmin^10^, which serve as an alternative antigen targets for cancer therapy^36^. In addition, glycoRNAs can engage P-selectin^7^ and Siglec family proteins^3, 23, 26^, thereby contributing to regulate neutrophil recruitment to inflammatory sites and cell-cell interaction^5, 7, 23, 26^. GlycoRNAs also form a molecular complex with heparan sulfate and regulate vascular endothelial growth factor A (VEGF-A) signaling during angiogenesis^11^. These findings indicate that the protein environments surrounding glycoRNAs can affect and determine their biological functions.

Despite this progress, current studies of glycoRNA function and protein associations have investigated glycoRNAs as pooled populations. As a result, it is largely unknown how the sequence identity of an individual glycoRNA is related to its surrounding proteins and biological function. Two central questions remain unresolved: 1) Which proteins interact with an individual, sequence-defined glycoRNA on a living-cell surface? 2) Will glycoRNAs with different sequences interact with the same set of proteins or do they have different protein interactome profiles? Answering these questions is essential for distinguishing biological functions that are broadly shared among glycoRNAs from the functions associated with specific glycoRNA species.

RNA-centric proximity-labeling technologies have recently provided a powerful framework for addressing such questions by coupling sequence-specific RNA recognition with proximity labeling to map proteins surrounding defined RNA species in their native cellular context^37–44^. However, these approaches were developed for conventional RNAs and do not address the unique molecular architecture of glycoRNAs, in which a glycan moiety is covalently linked to the RNA backbone. Till now, no proximity-labeling strategy has been developed to interrogate sequence-defined glycoRNAs in their native cell-surface environment. Existing approaches to glycoRNA-protein association are not well-suited to resolving this problem. For example, confocal colocalization imaging and flow cytometry are commonly used to evaluate the associations between glycoRNAs with selected candidate cell-surface proteins^3, 7, 10^. These approaches are valuable for targeted validation but have limited throughput. Proteome-wide discovery has been achieved by immobilizing chemically modified glycoRNAs on magnetic beads, incubating them with cell lysates and identifying enriched proteins by mass spectrometry^45^. However, chemical modification and immobilization of glycoRNAs may alter RNA folding or protein recognition, whereas cell lysis disrupts the native membrane environment and can change the organization and structures of membrane proteins. Most importantly, all existing approaches identify proteins associated with pooled glycoRNA populations rather than with one defined RNA sequence. Consequently, these methods do not readily enable sequence-resolved profiling of glycoRNA-associated proteins on intact cells (Fig. 1a). A novel approach is therefore needed to map the protein interactome of individual, sequence-defined glycoRNAs while preserving the organization of the living-cell surface (Fig. 1b).

**Fig. 1.**
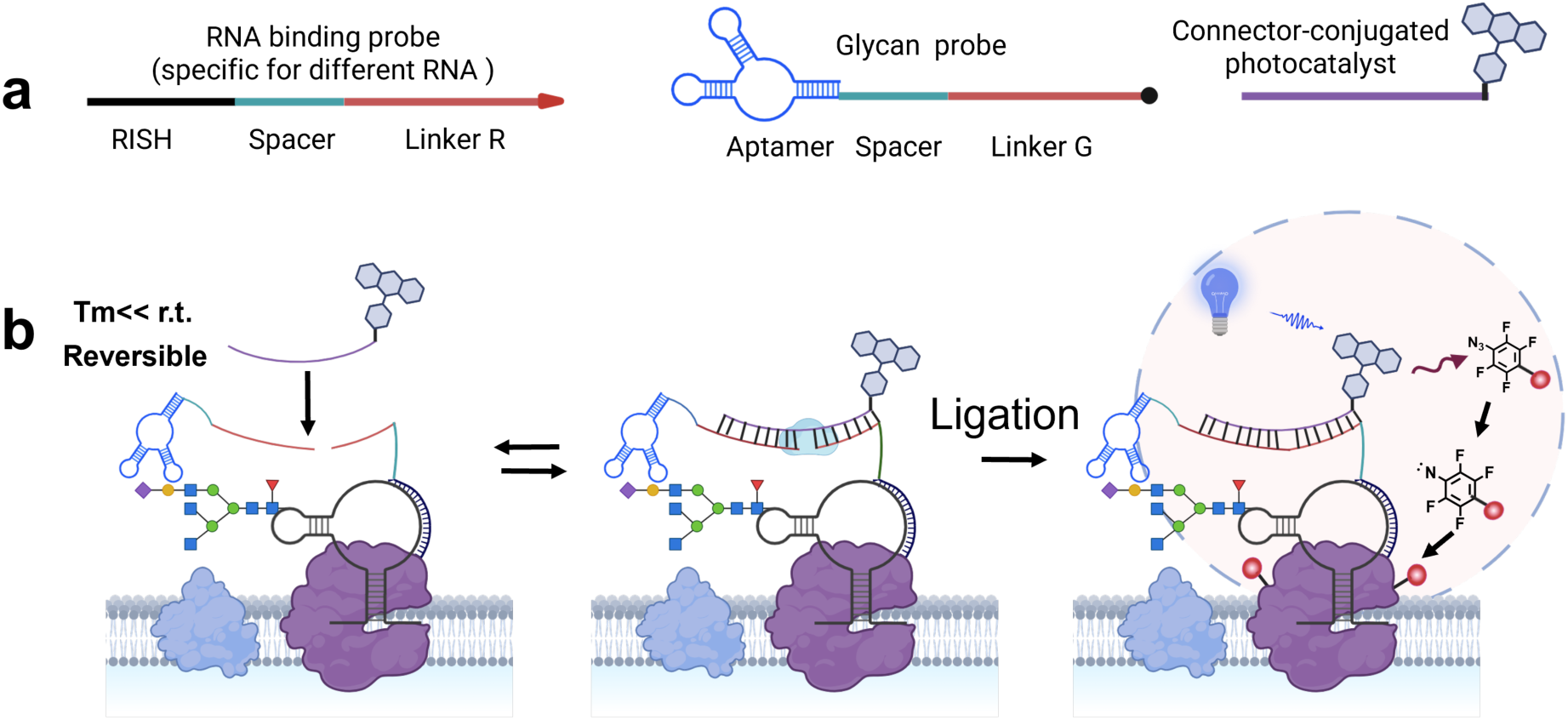
Programmable assembly of GlycoRNA-Map enables sequence-resolved *in situ* profiling of glycoRNA-protein interactomes. **a**, Design of the three-component GlycoRNA-Map system. An RNA-binding probe and a glycan probe recognize the RNA and glycan moieties of glycoRNA, respectively. Complementary linker regions on the two probes recruit a photocatalyst strand. **b**, Mechanism of *in situ* assembly and photocatalytic labeling. The recognition probes undergo reversible hybridization with the photocatalyst strand through the linker regions, followed by ligation to form a stable photocatalytic module on the target glycoRNA. Upon visible-light irradiation, the photocatalyst activates the photoactivatable labeling probe, enabling proximity labeling of proteins surrounding the target glycoRNA.

Herein, we develop an RNA sequence-selective proximity-labeling method, GlycoRNA-Map, to profile glycoRNA-protein interactome on living cell surfaces. This approach uses dual probes to recognize glycoRNAs and subsequently recruits a photocatalyst for localized protein labeling (Fig. 1c). To target glycoRNA with a specific sequence, we employed two probes (Fig. 1c): 1) A glycan-binding probe containing a sialic acid aptamer, a single-stranded nucleic acid designed to bind sialic acid in the glycan moiety of glycoRNAs. 2) an RNA *in situ* hybridization probe (RISH) attaching to the RNA moiety of the glycoRNAs. The co-recognition of glycoRNAs by these two probes can recruit the photocatalyst strand for proximity labeling. Upon light irradiation, this assembled complex can absorb photonic energy from visible light, transferring it to photolabeling probes (diazirine-biotin, tetrafluorophenyl azide-biotin (PhF_4_N_3_-biotin), phenyl azide-biotin (PhN_3_-biotin), or phenol-biotin) in the microenvironment, thereby labeling nearby glycoRNA-associated proteins with distinct effective labeling radii. Then, these labeled proteins can be isolated with streptavidin-coated magnetic beads and identified by mass spectrometry with label-free quantitation. By combining glycoRNA-sequence recognition and photocatalytic proximity labeling, the method maps the molecular environment of an individual glycoRNA without disrupting the plasma membrane through cell lysis. We applied the method to profile the protein interactome of five glycoRNAs, including Y5 RNA (Y5), SNORD1 (U1), SNORD3a (U3), SNORD8 (U8) or SNORD35a (U35a). These glycoRNAs were associated with both shared and unique sets of proximal proteins, indicating that individual glycoRNAs occupy partially distinct protein neighborhood environments (Fig. 1d). Notably, transforming growth factor-β receptors 1 and 2 (TGFBR1 and TGFBR2) were identified in the proximal proteomes of multiple glycoRNAs. Subsequent experiments showed that glycoRNA depletion by extracellular RNase treatment enhanced TGF-β receptor signaling and promoted cell migration, supporting a functional relationship between glycoRNAs and TGF-β signaling pathway (Fig. 1d). Finally, our system utilized commercially available fluorophore-modified oligonucleotides as programmable photocatalytic probes, which eliminate the need for genetic manipulation or customized conjugation and are broadly accessible for the scientific community. Together, our work establishes a general strategy for sequence-resolved profiling of glycoRNA-associated protein environments on living-cell surfaces. Our strategy extends RNA-centered proximity labeling beyond RNA identity to its chemical modification state on living-cell surfaces. This approach provides a general framework for investigating how chemically modified RNAs organize membrane signaling, cell-cell communication, and other biological processes at the surfaces of living cells.

## Results

### Development of a broadly accessible, multiscale RNA-centric proximity-labeling platform for glycoRNA

To establish a broadly accessible RNA-centric proximity-labeling platform, we repurpose commercially available fluorophore-modified oligonucleotides as programmable photocatalytic modules, which eliminate the need for customized enzyme or photocatalyst conjugation and genetic manipulation^46–51^. To design an ideal photocatalytic oligonucleotide, we selected carboxyfluorescein (FAM), carboxy-X-rhodamine (ROX) and tetramethylrhodamine (TAMRA) as candidate photocatalysts for platform development, as these fluorophores are widely commercial available and cost-effective oligonucleotide modifications and possess photochemical properties compatible with photocatalytic labeling^52–55^.

We first compared the photocatalytic activities of these candidate fluorophores using bovine serum albumin (BSA) as a model protein and diazirine-biotin as the photoactivatable labeling probe, with protein labeling efficiency assessed by western blot (WB) analysis (Fig. 2a). In the absence of a fluorophore, negligible protein labeling was detected under 470 nm or 520 nm irradiation condition, indicating that neither light irradiation nor diazirine-biotin alone was sufficient to induce proximity labeling (Fig. 2b,c). In contrast, all three fluorophores efficiently activated diazirine-biotin and labeled BSA under 470 nm laser irradiation (Fig. 2b,c), suggesting that fluorophore-modified oligonucleotides can function as photocatalytic modules. Among the three fluorophores, FAM consistently exhibited the highest labeling efficiency, approximately 1.25-fold and 1.52-fold compared with those of ROX and TAMRA, respectively (Fig. 2c). Given the visible-light absorption maxima of FAM, ROX and TAMRA (λ_max_ = 492, 578 and 550 nm, respectively; Supplementary Fig. 1), we further tested 520 nm excitation as a potentially more biocompatible alternative to blue light. All three fluorophores, including FAM, remained photocatalytically active (Fig. 2b,c). These results identified FAM as the optimal photocatalytic module for subsequent platform development.

**Fig. 2.**
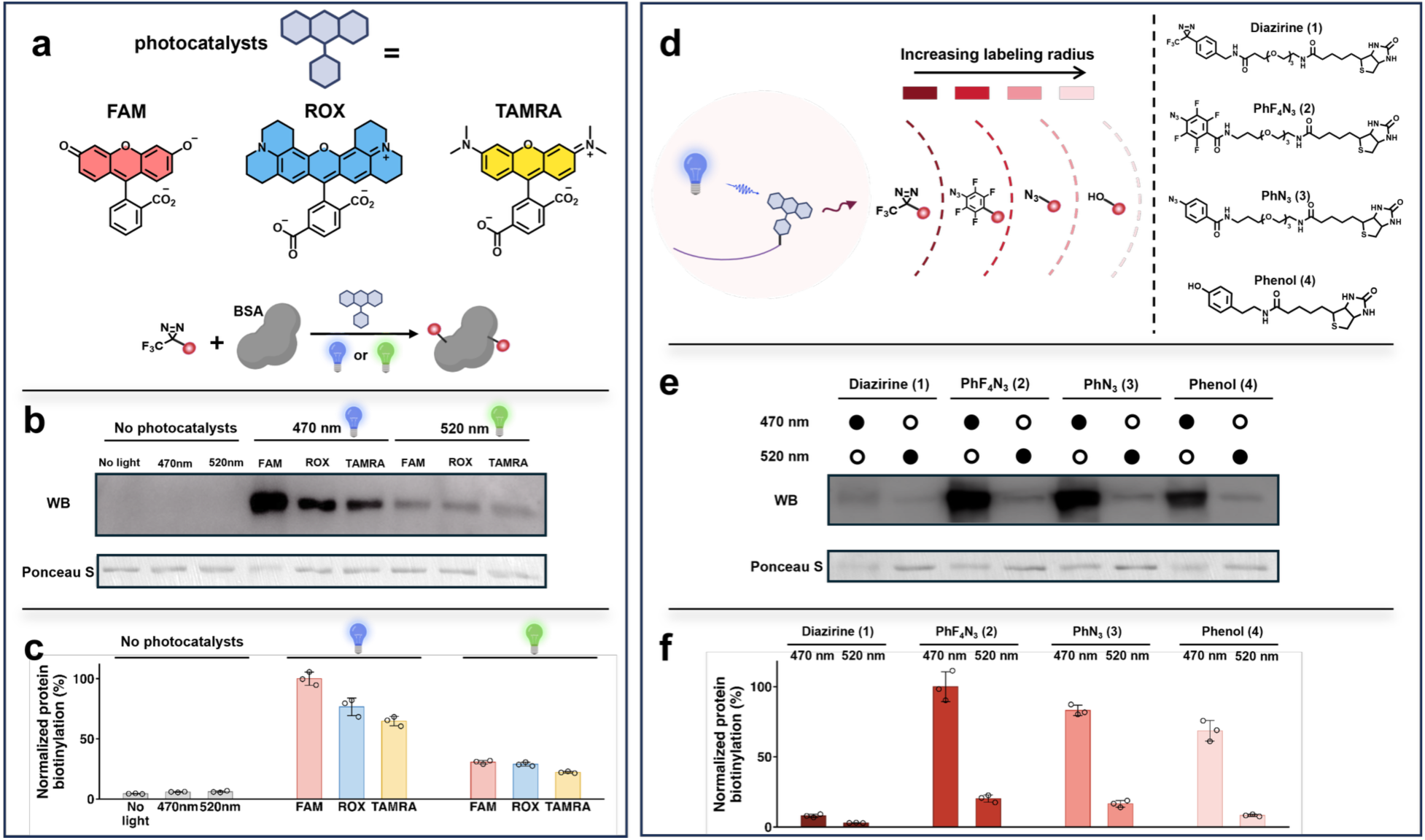
Development of an accessible, multiscale RNA-centric proximity-labeling platform for glycoRNA. **a,** Representative organic photocatalysts as commercially available oligonucleotide modifications: 6-carboxyfluorescein (FAM), 6-carboxy-X-rhodamine (ROX), and tetramethylrhodamine (TAMRA). **b,** Western blot (WB) showing fluorophore-mediated photocatalytic biotinylation of BSA. This experiment is performed with a diazirine-biotin photo-labeling probe under 470- or 520-nm LED illumination. **c,** Quantification of photocatalytic BSA biotinylation mediated by the three organic photocatalysts under different experimental conditions in (**b**). **d,** Schematic illustration of fluorophore-modified oligonucleotides coupled with photoactivatable labeling probes, including diazirine-biotin, tetrafluorophenyl azide-biotin (PhF_4_N_3_-biotin), phenyl azide-biotin (PhN_3_-biotin) and phenol-biotin, providing distinct effective labeling radii for the GlycoRNA-Map. **e,** WB analysis of FAM-mediated photocatalytic biotinylation of BSA using four photoactivatable labeling probes under 470-nm LED illumination. **f**, Quantification of the WB analysis shown in (**e**).

Because different photoactivatable labeling probes provide distinct effective labeling radii, compatibility with multiple probes is essential for establishing a versatile and fine-tunable proximity labeling platform to study RNA-centric protein interactome. We therefore evaluated four labeling probes, diazirine-biotin, PhF_4_N_3_-biotin, PhN_3_-biotin, and phenol-biotin (Fig. 2d), under both 470 nm and 520 nm irradiation. Using FAM as the photocatalyst, the four labeling probes exhibited markedly different labeling efficiencies. Under 470 nm irradiation, PhF_4_N_3_-biotin exhibited the highest labeling efficiency, whereas PhN_3_-biotin, phenol-biotin, and diazirine-biotin showed labeling efficiencies of ∼85%, ∼70%, and ∼8%, respectively, relative to PhF_4_N_3_-biotin. Under 520 nm irradiation, the corresponding labeling efficiencies were ∼20%, ∼16%, ∼8%, and ∼5%, respectively (Fig. 2e,f). Thus, FAM efficiently activated all four labeling probes, demonstrating broad compatibility across chemically distinct labeling probes while providing a range of labeling efficiencies. Among these probes, diazirine and aryl azides, including PhF_4_N_3_-biotin and PhN_3_-biotin, generate highly reactive carbenes or nitrenes with short lifetimes (∼2-10 ns), enabling relatively short-range protein labeling (∼10-70 nm) with broad amino acid coverage for nearest-neighborhood analysis. PhF_4_N_3_-biotin exhibits a shorter labeling radius than PhN_3_-biotin, likely owing to the higher reactivity of its fluorinated intermediates. In contrast, phenol-biotin generates longer-lived phenoxyl radicals (> 100 μs), supporting labeling distances of up to ∼300 nm. Consistent with these theoretical considerations, a super-resolution microscopy-based assay directly quantified proximity labeling radii by experiment, which reported 54 ± 12 nm for diazirine, 68 ± 17 nm for PhF_4_N_3_-biotin, 119 ± 33 nm for PhN_3_-biotin, and 269 ± 41 nm for phenol-biotin. By choosing different labeling probes, we can establish a modular and multiscale photocatalytic labeling strategy capable of interrogating glycoRNA-centered molecular microenvironments at different spatial scales. In this work, we will choose the labeling probe to achieve a balance between high labeling efficiency and precise labeling distance. Under identical irradiation conditions, PhF_4_N_3_-biotin consistently produced the strongest protein labeling efficacity, whereas diazirine-biotin exhibited the weakest activity (Fig. 2f). Especially, under 470 nm irradiation, PhF_4_N_3_-biotin generated approximately 11-fold stronger labeling than diazirine-biotin (Fig. 2f). Although diazirine-biotin provides the shortest labeling radius of approximately 50 nm, its labeling efficiency was substantially lower than that of PhF_4_N_3_-biotin, which retains a highly localized labeling radius of approximately 65 nm while providing markedly improved labeling efficiency. We therefore selected PhF_4_N_3_-biotin for subsequent glycoRNA-protein interactome profiling.

We next standardized the operating parameters of the proximity labeling platform by systematically optimizing the photocatalytic reaction conditions, including the concentrations of FAM and PhF_4_N_3_-biotin and the irradiation time. Protein labeling increased progressively with FAM concentration, demonstrating a clear fluorophore concentration dependence (Extended Data Fig. 1). Likewise, increasing the PhF_4_N_3_-biotin concentration enhanced protein labeling (Extended Data Fig. 2). In addition, time-dependent accumulation of biotinylated BSA was observed by WB analysis, with labeling reaching a plateau within 10 min under 470 nm illumination (Extended Data Fig. 3). These results demonstrate robust and tunable photocatalytic performance across a defined operation range. Based on these optimizations, FAM, PhF_4_N_3_-biotin, and 470 nm illumination were chosen as the standardized operating configuration for subsequent experiments.

To validate that the optimized platform can be directly implemented using commercially available fluorophore-modified oligonucleotides, we utilized a 5′-FAM-modified oligonucleotide targeting small nuclear glycoRNA U1 and U8 (U1-FAM or U8-FAM) and evaluated its photocatalytic performance under the optimized conditions. Using 100 μM PhF_4_N_3_-biotin and 470 nm irradiation for 10 min, the U1-FAM and U8-FAM probes efficiently mediated protein labeling in a probe concentration-dependent manner over the range of 1 to 10 μM (Extended Data Fig. 4). At a fixed U1-FAM concentration of 10 μM and PhF_4_N_3_-biotin concentration of 100 μM, labeling efficiency increased progressively with irradiation time under 470 nm illumination (Extended Data Fig. 5). These results demonstrate that commercially available FAM-modified oligonucleotide probes can be directly deployed as functional photocatalytic modules without additional chemical conjugation or post-synthetic modification.

Together, these studies establish a standardized, modular, and broadly accessible RNA-centric photocatalytic proximity labeling platform based on commercially available fluorophore-modified oligonucleotide probes. The platform provides broad accessibility, conjugation-free implementation, multiscale labeling capability, and RNA sequence programmability, representing a broadly adoptable platform for sequence-directed RNA-centric protein interactome.

### Programmable *in situ* assembly of glycoRNA-targeted photocatalytic probes on cells

To investigate glycoRNA-protein interactome on cell surface, we designed photocatalytic probes, named GlycoRNA-Map, which comprises three functional components (Fig. 1b and 3a): (1) a glycan-recognition probe containing a DNA aptamer that selectively binds N-acetylneuraminic acid (Neu5Ac), which is enriched in glycoRNAs, a spacer to minimize steric hindrance during hybridization, and a DNA linker (linker G) to attract photocatalyst strand; (2) an RNA-targeting probe containing RISH sequence, a spacer, and a DNA linker (linker R) to hybridize with photocatalyst strand; (3) a FAM-modified photocatalyst strand that hybridizes with linkers G and R to enable in situ ligation. At room temperature, the photocatalyst strand remains in a dynamic state of annealing with and detaching from the glycan-recognition probe and RNA-targeting probe. Only when these 3 probes are *in situ* assembled, T4 DNA ligase can ligate the glycan-recognition probe and the RNA-targeting probe and ensure their stable hybridization with the photocatalyst strand. The photocatalyst strand hybridizing to only one probe (either glycan-recognition probe or RNA-targeting probe) is then washed away. Therefore, the photocatalytic probe can be installed onto native cell surface glycoRNAs selectively and stably, and then serve as the photocatalyst for glycoRNA-centered proximity labeling (Fig. 1b and 3a).

**Fig. 3.**
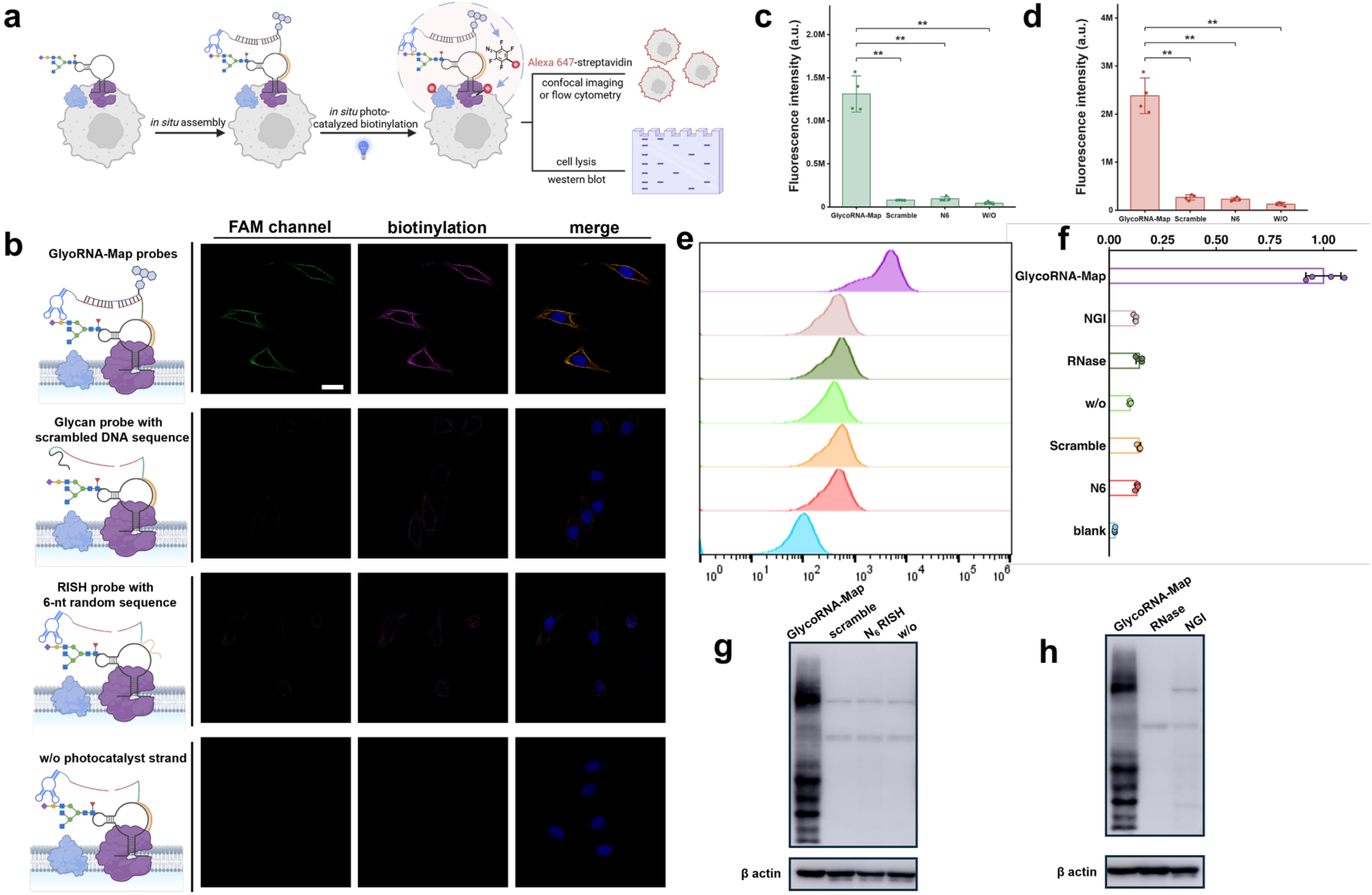
Programmable *in situ* assembly enables selective on-cell proximity labeling of glycoRNAs. **a**, Schematic illustration of on-cell labeling workflow using GlycoRNA-Map and detection of biotinylated proteins by streptavidin-fluorophore staining and WB analysis. **b**, Confocal microscopy images showing *in situ* assembly and on-cell biotinylation by GlycoRNA-Map in HeLa cells under various control conditions replacing individual components of GlycoRNA-Map. w/o, without. Scale bar, 20 μm. **c,d**, Quantification of FAM and streptavidin-Alexa 647 fluorescence intensities in **b** (n = 4). Significance was assessed by t-test. **P < 0.01, ***P < 0.001, ****P < 0.0001. **e**, Flow cytometric analysis of on-cell biotinylation mediated by GlycoRNA-Map. **f**, Quantification of flow cytometry data in (**e**) (n=4). Significance was assessed by t-test. **P < 0.01, ***P < 0.001, ****P < 0.0001. **g**, WB analysis of *in situ* assembly and on-cell biotinylation mediated by GlycoRNA-Map in HeLa cells under various control conditions, replacing individual components of GlycoRNA-Map. β-actin was used as a loading control. w/o, without. **h**, WB validation of the specificity of GlycoRNA-Map. Live HeLa cells were pretreated with RNase A/T1 or the glycosylation inhibitor NGI-1 before GlycoRNA-Map workflow.

We then applied GlycoRNA-Map to target the glycosylated small nuclear RNA U1 (U1 glycoRNA) as a representative cell-surface glycoRNA in HeLa cells. Following *in situ* assembly and photocatalytic labeling, cells were incubated with streptavidin-Alexa Fluor 647 to detect protein biotinylation generated by photocatalytic proximity labeling through confocal laser-scanning microscopy (CLSM) imaging (Fig. 3a). As shown in Fig. 3b, green fluorescence signal was observed on the cell surface, representing successful assembly of the FAM-modified photocatalyst strand to cell surface U1 glycoRNA. In addition, the existence of Alexa Fluor 647 (red signal) on cell membrane demonstrates that glycoRNA binding proteins are biotinylated through photocatalytical proximity labeling. Strong membrane-associated green and red fluorescence signals were observed only under conditions with all functional components of GlycoRNA-Map. Replacing the Neu5Ac aptamer with a scrambled aptamer, replacing the RNA RISH probe with a non-targeting random sequence (N6 probe), or omitting the FAM-modified photocatalyst strand reduced the corresponding green signals by 16.8-, 14.0- and 29.5-fold, respectively, and the red signals by 9.0-, 10.6- and 19.2-fold, respectively (Fig. 3c,d). These results demonstrate that selective photocatalytic labeling requires in situ assembly mediated by simultaneous recognition of both the RNA sequence and glycan moiety of cell surface glycoRNAs.

To determine whether GlycoRNA-Map specifically targets glycoRNAs, we manipulated glycoRNA abundance by independently perturbing the RNA and glycan components. To disrupt the RNA moiety of glycoRNAs, live cells were treated with RNase A (0.2 mg/mL) and RNase T1 (200U/mL) at 25 °C for 20 min before imaging. RNase treatment reduced the green fluorescence signal from FAM-modified photocatalyst strand by 90% and the corresponding cell-surface biotinylation signal by 88% relative to untreated cells (Extended Data Fig. 6), demonstrating that cell surface glycoRNA is required for efficient probe assembly and subsequent photocatalytic labeling. We next disrupted glycoRNA biosynthesis by treating cells with the N-linked glycosylation inhibitor NGI-1 at the concentration of 8 µM for 24 h. NGI-1 treatment reduced the fluorescence signal of FAM-modified photocatalyst strand (Green) and cell surface biotinylation (Red) by 86% and 90%, respectively (Extended Data Fig. 6). Together, these results demonstrate that both the RNA and glycan components of glycoRNAs are required for efficient GlycoRNA-Map-mediated photocatalytic proximity labeling.

The photocatalytic labeling across the cell population was further validated by flow cytometry using streptavidin-Alexa Fluor 647 fluorescence as the readout. Cells labeled by all functional components of GlycoRNA-Map exhibited significant fluorescence intensity, whereas replacement of the glycan-recognition aptamer with a scrambled aptamer, replacement of the RNA RISH probe with the N6 probe, omission of the photocatalyst strand, or disruption of glycoRNAs abundance by RNase or NGI treatment reduced fluorescence signal of streptavidin-Alexa Fluor 647 (Fig. 3e,f). These results were consistent with the labeling efficiencies observed by confocal imaging and further demonstrated efficient and selective glycoRNA-directed photocatalytic labeling at the single-cell level.

We next established a biochemical workflow to profile the glycoRNA-centered protein neighborhood by on-cell photocatalytic labeling followed by WB analysis (Fig. 3a). Hela cells are treated by GlycoRNA-Map workflow to perform on-cell photocatalytic labeling, and subsequently lysed to extract total proteins and then processed for selective staining of biotinylated proteins. As shown in Fig. 3g, robust protein biotinylation was detected by WB only after successful assembly of all 3 GlycoRNA-Map probes. In contrast, replacing the RNA RISH probe with the N6 probe, replacing the Neu5Ac-binding aptamer with a scrambled DNA sequence, or omitting the FAM-modified photocatalyst strand almost completely abolished protein labeling. RNase and NGI treatment markedly reduced protein labeling relative to untreated cells (Fig. 3h), demonstrating that efficient photocatalytic labeling requires intact cell-surface glycoRNAs containing both the target RNA sequence and a glycan moiety. Together, these results establish that GlycoRNA-Map enables programmable *in situ* assembly of a photocatalyst strand onto cell-surface glycoRNAs through dual recognition of their RNA sequence and glycan moiety. The assembled modules can then directly mediate *in situ* proximity labeling, providing a sequence-programmable strategy for profiling glycoRNA-centered protein neighborhoods.

### Sequence-resolved glycoRNA-protein interactome reveals distinct glycoRNA-centered protein neighborhoods

Having established a robust platform for glycoRNA-centered photocatalytic labeling, we next investigated how protein neighborhoods surrounding cell-surface glycoRNAs vary with RNA sequence. To address this question, we designed RISH sections in the RNA-binding probes to be replaced with different sequences targeting individual glycoRNAs, such as U1, U3, U8, U35a or Y5 (Fig. 4a). As such, these RNA specific GlycoRNA-Map platforms can be utilized to profile the corresponding glycoRNA-protein interactome with RNA sequence resolution in the human lung adenocarcinoma epithelial cell line, A549 cells (Fig. 4a). To serve as a negative control group, the RNA-binding probe with a randomized probe library of 6 nt oligonucleotides (N6 probe) was utilized, which cannot hybridize with any glycoRNAs or support *in situ* assembly of the photocatalytic module. In addition, tetrafluorophenyl azide was chosen as the photocatalytic labeling probe. Cells were treated with GlycoRNA-Map with different RNA RISH probes (U1, U3, U8, U35a, Y5, and N6), following photocatalytic labeling. Thereafter, biotinylated proteins were enriched on streptavidin-functionalized beads and digested on-bead with trypsin. Samples were prepared in biological duplicates for MS analysis by using label-free quantitation. By comparing each glycoRNA proximal proteome with the N6 control, we identified 200 significantly enriched proteins for U1, 229 for U3, 418 for U8, 258 for U35a, and 262 for Y5, using thresholds of log₂(fold change) > 1 and P < 0.05 (Fig. 4b). These results demonstrate robust sequence-directed mapping of glycoRNA-centered protein neighborhoods. The full list of identified proteins is provided in Table S1. Notably, 64% of the 845 significantly enriched proteins were detected exclusively in a single glycoRNA dataset and were absent from the other four datasets. The remaining proteins showed limited overlap across glycoRNA datasets: 19.4% (164 proteins) were detected in two datasets, 9.3% (79 proteins) in three, 5.2% (44 proteins) in four, and only 2.0% (17 proteins) across all five datasets. These results indicate that different glycoRNA sequences are associated with highly distinct protein neighborhoods (Fig. 4b,c,d).

**Fig. 4.**
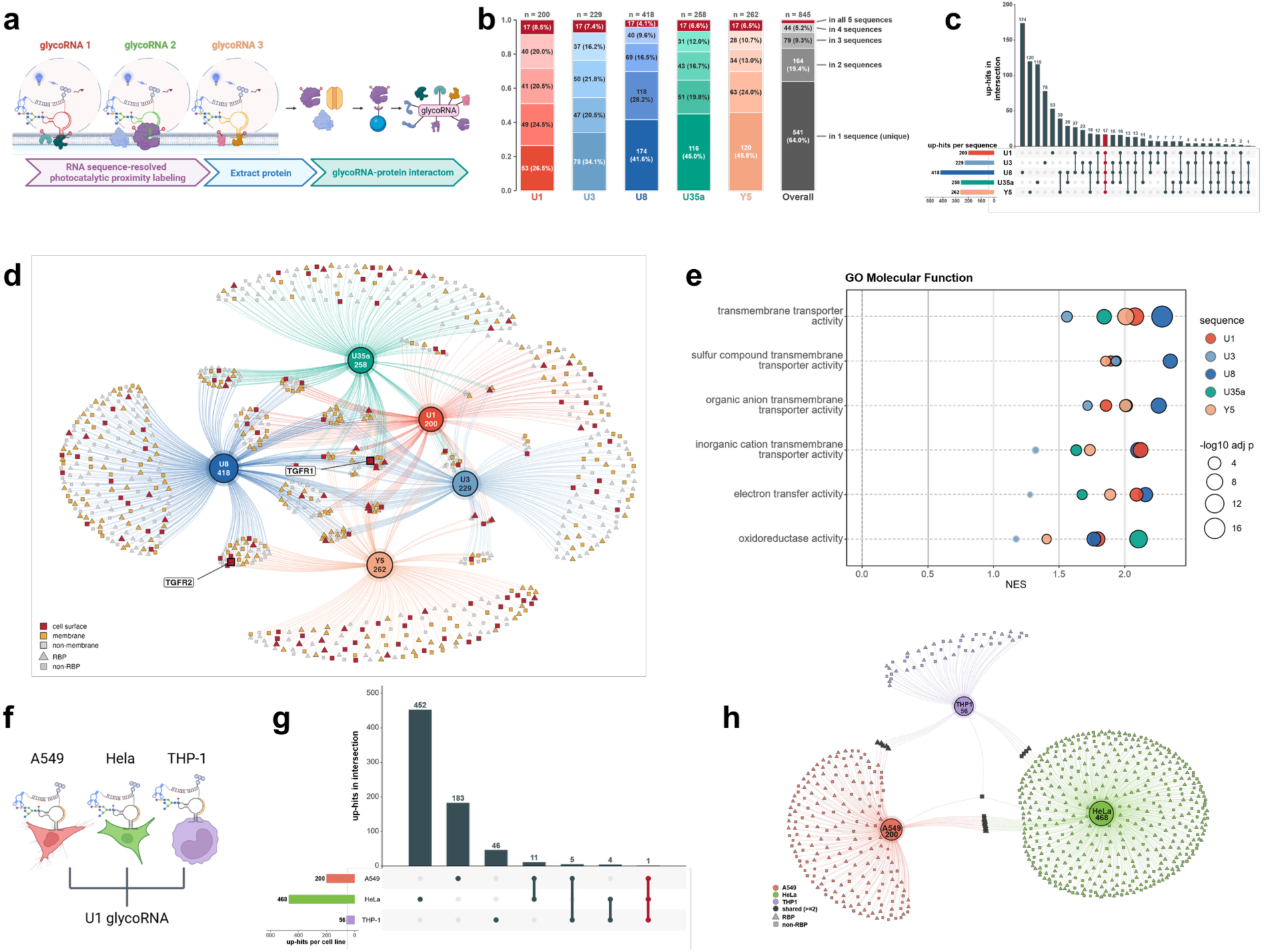
Sequence and cellular context shape glycoRNA-protein interactomes. **a**, Schematic illustration of sequence-resolved glycoRNA-protein interactome profiled by GlycoRNA-Map. **b**, Overlap of significantly enriched proteins across U1, U3, U8, U35a and Y5 glycoRNA datasets. Stacked bars indicate proteins detected uniquely or shared among two, three, four or all five datasets. **c**, UpSet plot showing the overlap of significantly enriched proteins across the five glycoRNA datasets. **d**, Network visualization of protein neighborhoods associated with U1, U3, U8, U35a and Y5 glycoRNAs. TGFBR1 and TGFBR2 are highlighted. **e**, Gene Ontology (GO) molecular function enrichment analysis of the five glycoRNA-associated protein neighborhoods. Dot position indicates the normalized enrichment score, dot size indicates −log10 adjusted P value, and dot color denotes the corresponding glycoRNA. **f**, Schematic illustration of cross-cell-type profiling of the U1 glycoRNA-associated protein neighborhood in A549, HeLa and THP-1 cells using GlycoRNA-Map. **g**, UpSet plot showing the overlap of significantly enriched proteins associated with U1 glycoRNA across A549, HeLa and THP-1 cells. **h**, Network visualization of U1 glycoRNA-associated protein neighborhoods across A549, HeLa and THP-1 cells.

Gene Ontology (GO) cellular component (CC) analysis showed that 47.6% of the enriched proteins were primarily associated with membranes and the cell surface, supporting the specificity of our labeling method (Extended Data Fig. 7). Gene Ontology (GO) analysis of biological process and molecular function terms revealed strong enrichment for functions related to transmembrane transporter activity (Fig. 4e). Several highly enriched membrane proteins, including SLC1A4, SLC29A1 and TGFBR1, are involved in amino acid and nucleoside transport or the transduction of extracellular signals across the plasma membrane. Similar to previous findings that glycoRNA-csRBP clusters facilitate cell-penetrating peptide entry, our results further suggest potential roles for glycoRNA-associated protein assemblies in organizing functional microenvironments at the cell surface.

Previous studies have shown that a subset of RNA-binding proteins (RBPs) resides on the cell surface, where cell-surface RBPs (csRBPs) form defined nanoclusters in proximity with glycoRNAs. We next assessed the RNA-binding potential of the enriched proteins using RBP2GO, a database of validated and putative RNA-binding proteins (RBPs). Among the 845 enriched proteins in all 5 glycoRNA datasets, 372 (44.0%) were annotated as RBPs. For each glycoRNA datasets, RBP proportions were found to be 40.0%, 49.3%, 45.9%, 43.4% and 40.5% for U1, U3, U8, U35a and Y5 glycoRNAs, respectively (Extended Data Fig. 8). All 372 proteins had an RBP2GO score ≥5, corresponding to high-confidence RBP annotations. Compared with another independent RBP database generated by Perr *et. al.*, we identified 393 proteins (46.5%) among our significantly enriched proteins, including nucleolin (NCL) and the DNA-dependent protein kinase catalytic subunit (DNA-PKcs), both of which are among the top-ranked proteins in this RBP database (Extended Data Fig. 8). Together with previous reports showing that RBPs can localize on the plasma membrane and form nanoclusters in proximity to glycoRNAs, our findings support a substantial RBP as glycoRNA binders on cell membranes.

Furthermore, we asked whether glycoRNA-centered protein neighborhoods are conserved across different cell types, utilizing U1 glycoRNA as an example (Fig. 4f). Using RNA-binding probe targeting the U1, GlycoRNA-Map profiled the proximal proteomes of U1 glycoRNA in HeLa, A549 and THP-1 cells, identifying 468, 200 and 56 significantly enriched proteins, respectively (Fig. 4g). Despite targeting the identical RNA sequence, U1 glycoRNA exhibited strikingly distinct protein neighborhoods across the three cell lines, with minimal overlap among the enriched proteins. Only one enriched protein was shared across all three cell lines, while 20 proteins were detected in two cell types. Collectively, the shared proteins, including those detected in either two or three cell lines, accounted for only 3.0% of the total enriched proteins, whereas the vast majority were uniquely detected in individual cell lines (Fig. 4h). These results indicate that glycoRNA-protein interactome are shaped not only by RNA sequence but also by cellular context.

### Sequence-directed interactomes identifies glycoRNA-regulated TGFβ receptor microenvironment

The enrichment of membrane proteins involved in transmembrane transport and signaling within glycoRNA-proximal proteomes inspired us to investigate whether these proteins are functionally regulated by cell-surface glycoRNAs. Among the numerous membrane proteins identified in our datasets, TGF-β receptor 1 (TGFBR1) was consistently among the top-enriched proteins across all five glycoRNA sequences, including U1, U3, U8, U35a, and Y5. In addition, TGF-β receptor 2 (TGFBR2) was selectively enriched within the U8 and Y5 glycoRNA neighborhoods (Fig. 5a and Extended Data Fig. 9). Given the central role of the TGF-β receptor complex in regulating cell migration and other extracellular signaling processes, we selected this receptor system for functional investigation. First, it is important to determine and cross-validate whether glycoRNAs are spatially associated with TGF-β receptors at the cell surface. To answer this question, we examined the spatial distribution of glycoRNAs and TGFBR1/2 by confocal imaging. TGFBR1/2 were detected by immunofluorescence (red), while individual glycoRNAs were visualized by RNA in situ hybridization (RISH; green). As shown in Fig. 5b and Extended Data Fig. 10, TGFBR1 was densely localized at the plasma membrane of A549 cells. RISH targeting U1, U8, and U35a glycoRNAs produced promising cell-surface signals, which showed clear spatial colocalization with TGFBR1. Pearson’s correlation coefficients between glycoRNAs and TGFBR1 were 0.75 for U1, 0.86 for U8, and 0.79 for U35a. Similarly, U8 glycoRNA showed substantial colocalization with TGFBR2, with a Pearson’s correlation coefficient of 0.78. These results support the close spatial association between glycoRNAs and TGF-β receptors, which occupy a shared cell-surface microenvironment. In addition to the biochemical validation of the interaction between glycoRNA and TGFBR1/2, we complemented our analysis with AlphaFold 3 (AF3), which uses artificial intelligence to generate plausible models of RNA-protein complexes (Extended Data Fig. 11).

**Fig. 5.**
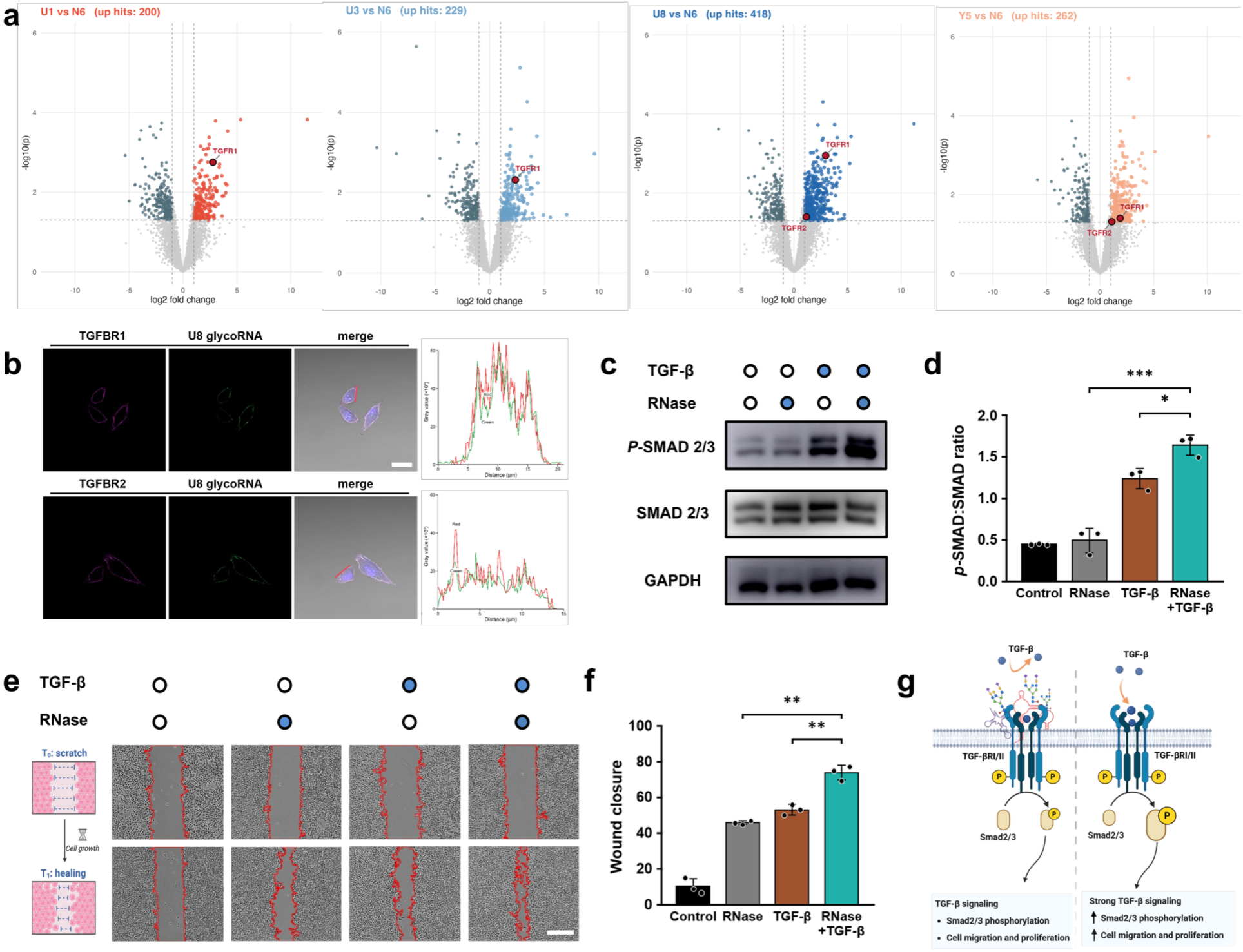
GlycoRNA regulates TGF-β receptor signaling pathway. **a**, Volcano plots showing significantly enriched proteins in the U1, U3, U8 and Y5 glycoRNA-associated proteins relative to the N6 control. TGFBR1 and TGFBR2 are highlighted. The x axis represents log2 fold change and the y axis represents −log10(P value). **b**, Confocal microscopy showing the spatial association of U8 glycoRNA with TGFBR1 and TGFBR2 at the cell surface. U8 glycoRNA was visualized by RNA *in situ* hybridization (green), whereas TGFBR1 and TGFBR2 were detected by immunofluorescence (magenta). Scale bar, 20 μm. Merged images and fluorescence intensity profiles along the indicated lines are shown. **c**, Western blot analysis of phosphorylated SMAD2/3 (p-SMAD2/3) and total SMAD2/3 in A549 cells treated with TGF-β1, RNase A/T1, or both. Untreated cells were used as the control. GAPDH was used as a loading control. Scale bar, 500 μm. **d**, Quantification of p-SMAD2/3 levels normalized to total SMAD2/3 in c. **e**, Representative images of wound-healing assays in A549 cells under the indicated treatment conditions. Cells were treated with TGF-β1, RNase A/T1, or both, and wound closure was monitored over time. **f**, Quantification of wound closure in e. **g**, Proposed model illustrating the role of cell-surface glycoRNAs in modulating TGF-β receptor signaling. Cell-surface glycoRNAs spatially associate with the TGF-β receptor complex and restrict ligand accessibility, whereas removal of cell-surface glycoRNAs potentiates TGF-β-induced SMAD2/3 phosphorylation and downstream cell migration and proliferation. n.s., not significant; **P < 0.01, ***P < 0.001, ****P < 0.0001.

We next asked whether the spatial proximity of glycoRNAs to TGFBR1/2 reflects a functional interaction that modulates the canonical TGF-β signaling pathway. We therefore considered three potential modes of glycoRNAs’ function: 1) glycoRNAs physically impede ligand binding or receptor assembly through steric or electrostatic effects; 2) glycoRNAs facilitate TGF-β engagement with TGFBR2 by acting as a co-receptor or promote TGFBR1/2 assembly into signaling-competent receptor complexes; 3) glycoRNAs function as bystanders without substantially affecting ligand binding or receptor assembly.

To determine whether glycoRNA can affect TGF-β signaling, we examined the activation of canonical SMAD pathway. As shown in Fig 5c,d, RNase A/T1 treatment alone did not induce detectable SMAD2/3 phosphorylation, indicating that removal of cell-surface glycoRNAs is insufficient to activate the pathway in the absence of TGF-β ligand. As expected, TGF-β1 stimulation increased phospho-SMAD2/3 levels relative to untreated cells. Remarkably, RNase A/T1 pretreatment further potentiated ligand-induced signaling, resulting in an approximately 2.2-fold increase in phospho-SMAD2/3 compared with TGF-β1 stimulation alone. These findings demonstrate that removal of the cell-surface glycoRNA enhances TGF-β-dependent receptor activation rather than directly activating the signaling pathway by itself.

Because activation of the TGF-β pathway promotes cell migration, we next performed wound-healing assays in A549 cells to investigate whether glycoRNAs affect cell migration, under four experimental conditions: untreated control, TGF-β1 stimulation alone, RNase A/T1 treatment alone, and RNase pretreatment followed by TGF-β1 stimulation. Consistent with the established role of TGF-β signaling, TGF-β1 significantly accelerated wound closure compared with untreated cells (Fig. 5e). RNase treatment alone also enhanced cell migration. Notably, RNase pretreatment followed by TGF-β1 stimulation produced the greatest migratory response, exceeding TGF-β1 treatment alone by approximately **2.2-fold** and RNase treatment alone by approximately **2.5-fold** (Fig. 5f). These findings suggest that cell-surface glycoRNAs inhibit the activation of TGF-β signaling pathway.

Collectively, these results support a model in which cell-surface glycoRNAs interact with TGF-β receptor to restrict receptor accessibility to extracellular ligands. Removal of the glycoRNAs increases ligand access to the receptor complex, thereby enhancing downstream SMAD signaling and promoting cell migration (Fig. 5g). Together with the RNA sequence-directed interactome analyses, these findings establish that glycoRNAs can functionally regulate extracellular signaling by interacting with their protein neighborhoods.

## Discussion

In this work, we developed a programmable photocatalytic proximity-labeling platform, termed GlycoRNA-Map, to map glycoRNA-centered protein neighborhoods at the cell surface. GlycoRNA-Map uses a glycan-binding aptamer and an RNA hybridization probe for dual recognition of the glycan and RNA moieties of glycoRNAs, respectively, followed by in situ assembly of a FAM-modified oligonucleotide that installs photocatalytic activity onto the target glycoRNA. The assembled photocatalytic module activates photoactivatable labeling probes upon light irradiation, enabling proteins proximal to glycoRNAs to be biotinylated, enriched and identified by mass spectrometry. Consequently, GlycoRNA-Map enables RNA sequence-directed profiling of glycoRNA-centered protein neighborhoods and reveals distinct protein microenvironments associated with different glycoRNA sequences and cellular contexts. GlycoRNA-Map has several advantages: (1) it uses commercially available fluorophore-modified oligonucleotides as photocatalytic modules, eliminating the need for customized enzyme or photocatalyst conjugation and making the platform broadly accessible for the scientific community; (2) GlycoRNA-Map employs dual recognition of the RNA and glycan moieties, enabling selective targeting of endogenous glycoRNAs rather than the glycan or RNA component alone; (3) the RNA recognition sequence can be readily reprogrammed, allowing customizable profiling of different glycoRNA sequences; and (4) GlycoRNA-Map provides a multiscale framework for resolving glycoRNA-associated protein neighborhoods, from spatially resolved cell-surface labeling to sequence-resolved proteomic profiling?

We next applied GlycoRNA-Map to investigate the organization and functional significance of cell-surface glycoRNA-associated protein neighborhoods. By profiling five glycoRNAs with distinct RNA sequences, we revealed that glycoRNA sequence is strongly associated with the composition of their proximal protein neighborhoods, and that the majority of enriched proteins are uniquely associated with individual glycoRNAs. We further examined the same U1 glycoRNA across different cell lines and found strikingly distinct protein neighborhoods in HeLa, A549 and THP-1 cells, indicating that glycoRNA-associated protein organization is shaped not only by RNA sequence but also by cellular context. GO analysis revealed strong enrichment of membrane proteins involved in transmembrane transport and extracellular signal transduction. Because TGFBR1 was consistently enriched across diverse glycoRNA-proximal proteomes, while TGFBR2 was enriched in the U8 and Y5 glycoRNA-proximal proteomes, we further examined their spatial association with glycoRNAs by confocal imaging and found clear colocalization of glycoRNAs with TGFBR1/2 at the cell surface. This observation led us to investigate the function of glycoRNA in regulating TGF-β signaling pathway. Removal of cell-surface glycoRNAs potentiates TGF-β-induced SMAD2/3 activation and cell migration. Together, these findings establish a functional connection between glycoRNA-protein interaction and extracellular signal transduction, and suggest that glycoRNAs’ function as regulator for TGF-β signaling pathway.

This work demonstrates that GlycoRNA-Map enables programmable, accessible profiling of glycoRNA-centered protein neighborhoods at the cell surface with RNA-sequence specificity. The modular design of GlycoRNA-Map provides a general framework for coupling programmable molecular recognition to photocatalytic labeling, enabling interrogation of the molecular environments surrounding defined RNA species. In addition, GlycoRNA-Map represents a powerful method to discover and investigate other potential functions and roles of glycoRNAs in many biological processes, including cell communication, host-immune response, and, in the process, uncover their relevance to diseases, such as cancer, autoimmune diseases, and inflammatory conditions. In the future, the photocatalytic module of GlycoRNA-Map could be integrated with single-molecule fluorescence in situ hybridization (smFISH) or multiplexed error-robust fluorescence in situ hybridization (MERFISH) to achieve spatially resolved RNA-centric proximity labeling with transcript-level or multiplexed RNA specificity, thereby extending the application of GlycoRNA-Map from the cell surface RNA to intracellular RNA. In parallel, replacing the recognition modules could enable this framework to target specific RNA modifications, such as m6A or m5C modified RNAs, expanding its utility as a versatile platform for programmable proximity labeling across diverse biomolecular contexts.

