## Supplementary file for "Programmable sequence-resolved *in situ* profiling of glycoRNA-protein interactomes by GlycoRNA-Map"

**Text S1: AlphaFold 3 predictions of binary RNA-protein complexes**

Based on AlphaFold 3 (AF3) docking predictions, the interaction between U1 and its target proteins involves one predicted binding site driven by a dual mechanism of hydrogen bonding and surface electrostatic complementarity (Figure S1A). In the hydrogen bond-dominated Binding Site 1, RNAs exhibit comparable interaction strengths with Transforming Growth Factor-beta Receptor 1 (TGF-β R1) and Transforming Growth Factor-beta Receptor 2 (TGF-β R2) (Table S1), targeting a densely positively charged protein surface (blue region). Binding Site 1 features a prominent positively charged patch (blue region) that exerts strong electrostatic attraction on the negatively charged RNA phosphate backbone; such non-specific Coulombic forces likely act synergistically during initial RNA capture. This feature is consistently reflected across other RNA–protein binding models (Figures S1–S2).

Across the RNA complexes, the two transforming growth factor-beta receptors show similar binding profiles. The RNA–TGF-β R1/2 models exhibit pTM values generally ranging between 0.5 and 0.6, indicating reasonable reliability in capturing global topology. Additionally, the U35a–TGF-β R1 interface shows an interface predicted TM-score (ipTM) of 0.42, reflecting moderate confidence in the predicted interface residue interactions (Table S1). Detailed docking analysis using PyMOL revealed that, except for U3 and Y5, the number of hydrogen bonds across the other docking models is roughly equivalent, indicating that the binding extent of the remaining RNAs to both proteins is comparable. Specifically, U3 exclusively favors binding to TGF-β R1, with no hydrogen bonding sites present for TGF-β R2. Conversely, Y5 tends to bind to TGF-β R2, exhibiting weaker binding to TGF-β R1. Nevertheless, a degree of uncertainty remains in the predictions generated by the AF3 model.

**Text S2: AlphaFold 3 predictions of the TGF-β R1–RNA–TGF-β R2 ternary complex**

As homologous proteins, TGF-β R1 and TGF-β R2 frequently act in concert within the cellular environment, rendering the investigation of their ternary complexes with RNA highly biologically relevant. In this study, three RNA–protein models with higher pTM scores (above 0.50) were selected for further binding predictions. Compared to the binary RNA–protein models, the predicted TGF-β R1–TGF-β R2 interaction model exhibits higher interface predicted TM-scores (ipTM) and predicted TM-scores (pTM), both approaching 0.60 (Table S1). This signifies a moderate level of confidence in the predicted residue-interface interactions, offering enhanced reliability over isolated binary counterparts. This interface is stabilized by 19 hydrogen bonds, corresponding to 19 interacting residue pairs (Figure S3A). The introduction of RNA molecules (U8, U35a, and Y5) results in a marginal decrease in ipTM, suggesting that the presence of RNA might sterically or energetically hinder protein oligomerization (Table S1). Notably, across all three ternary models, the number of hydrogen bonds stabilizing the TGF-β R1–TGF-β R2 interface remains largely unaltered, whereas the hydrogen-bonding binding sites between RNA and the TGF proteins are substantially increased compared to their isolated binary states.

The ternary models for U8 and U35a exhibit a consistent trend wherein the RNA–TGF-β R1 interaction predominates, with the number of hydrogen bonds far exceeding that of the RNA–TGF-β R2 interface. In contrast, the Y5 ternary binding model displays the opposite pattern, dominated by the Y5–TGF-β R2 interaction. Crucially, these RNA–TGF-β R associations occur without significantly disrupting the pre-existing hydrogen-bonding network between TGF-β R1 and TGF-β R2. These predicted outcomes align with the conclusions in Text S1, confirming that U8 and U35a bind more strongly to TGF-β R1 than to TGF-β R2, whereas Y5 exhibits the reverse selectivity. This shift in binding profiles implies that within the ternary assembly, TGF-β R2 interacts indirectly with RNA (U8 and U35a), primarily by forming a protein–protein interaction (PPI) dimeric network anchored by TGF-β R1 (Figure S3B–C). Conversely, for Y5, TGF-β R1 engages in an indirect interaction (Figure S3D). Distinct nucleic acids binding to the same protein model establish divergent interaction networks; specifically, homologous RNAs (U8 and U35a) display identical TGF-β Receptor binding preferences, contrasting sharply with Y5. Furthermore, the discrepancies between the predicted RNA–TGF-β R interfaces in ternary versus binary models underscore that nucleic acid–protein interactions establish distinct structural networks depending on their spatial conformations. The interaction landscape within a multimeric, higher-order conformation diverges sharply from that in an isolated binary state, thereby rendering the ternary model a more faithful representation of the crowded, complex macromolecular environment in living cells.

Through intricate cooperative effects, the ternary complex model delineates a more robust binding architecture than its binary counterparts. It elucidates a potential residue-level mechanism for the concurrent engagement of RNA with both homoproteins: TGF-β R2 forms a dimeric scaffold anchored to the TGF-β R1 interface while TGF-β R1 concurrently establishes tight associations with RNA (U8 and U35a) via robust hydrogen-bonding networks. Conversely, Y5 operates through a reverse binding mode centered on TGF-β R2 as the direct target. Further analysis reveals that the addition of RNA consistently leads to slight decreases in ipTM scores across all binding models (Table S1). This trend likely stems from inherent limitations in AlphaFold 3 predictions, specifically the pronounced conformational flexibility of RNA coupled with the relatively sparse representation of RNA structures in current training databases.

**Text S3: AlphaFold 3 predictions of the TGF-β R1–RNA–TGF-β R2 ternary complex**

Transforming growth factor-beta (TGF-β) regulates cellular metabolism by recruiting transforming growth factor-beta receptor 2 (TGF-β R2). To elucidate the impact of protein-binding RNAs (U8, U35a, and Y5) on this recruitment pathway, we performed structural simulations using AF3 (Table S2). During the recruitment of TGF-β R2 by TGF-β, the two proteins establish 17 pairs of interprotein interactions across 17 distinct anchoring sites (Figure S4 A). Upon the introduction of RNA, a consistent binding trend is observed: the RNA selectively associates with TGF-β rather than TGF-β R2 (Figure S3 B–D). This implies that, within these predictive models, all potential RNA-binding sites on TGF-β R2 are entirely occupied by TGF-β. Consequently, the recruitment capacity of TGF-β outcompetes the RNA–TGF-β R2 interactions, even surpassing the binding affinity of Y5 toward TGF-β R2. However, the addition of U35a nearly halves the ipTM of the complex to 0.18, compared to both the other RNA-containing ternary model and the protein–protein interaction model (about 0.36). This pronounced decrease indicates that the association of U35a with the protein complex is highly unstable, implying that U35a sterically or energetically perturbs the TGF-β recruitment process.

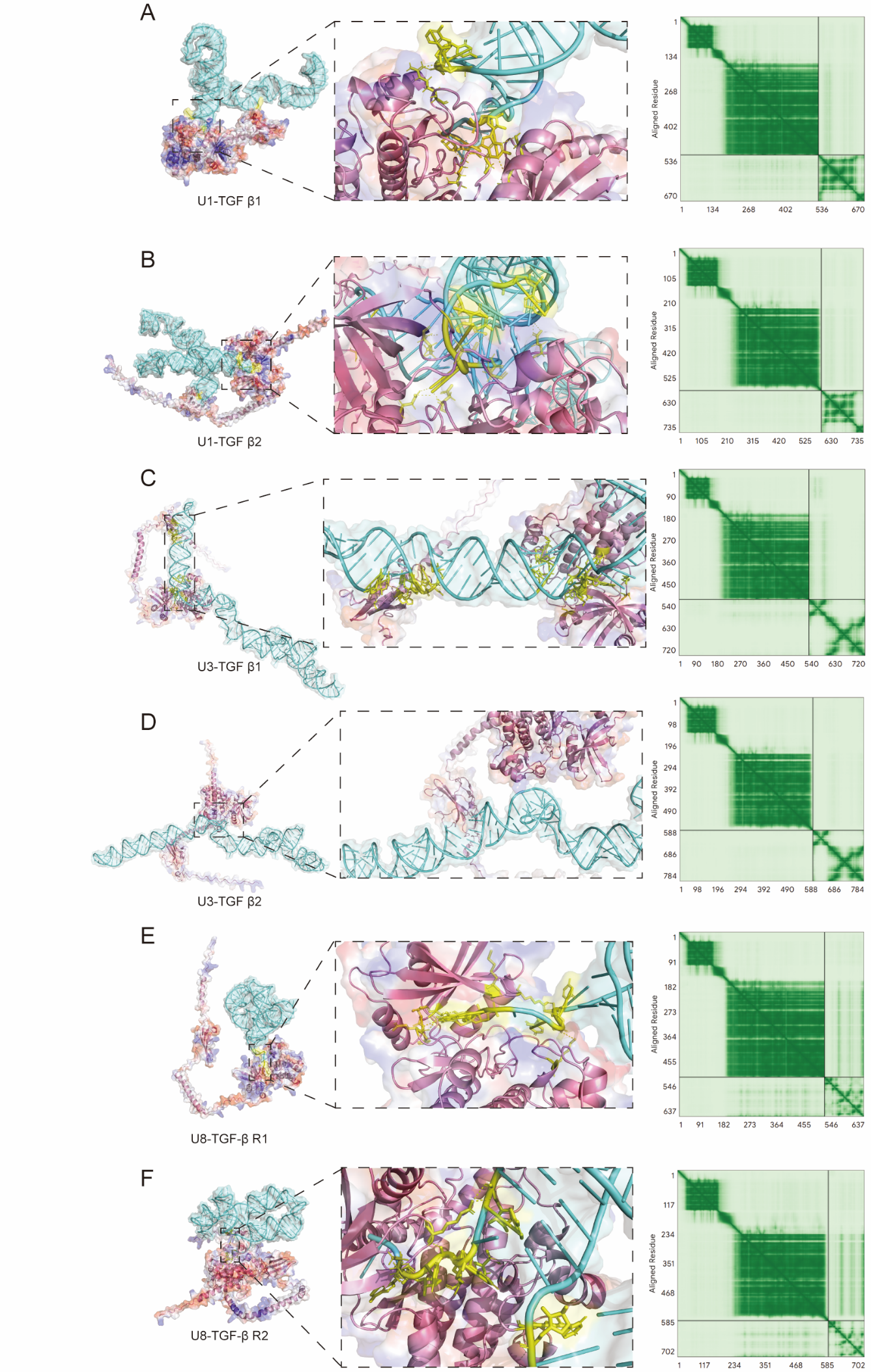

**Figure S1. AlphaFold 3 predictions of RNA–protein complexes.** Proteins and RNA are rendered as pink and cyan ribbons, respectively. Interacting amino acid residues and RNA bases are highlighted as yellow groups, with hydrogen bonds indicated by yellow dashed lines. On the electrostatic potential surface, blue and red patches represent densely positively and negatively charged regions, respectively. **A-B** Predicted U1–protein (TGF-β R1/2) complex model showing a single binding site, alongside the corresponding predicted alignment error (PAE) matrix. **C-D** Predicted U3–protein (TGF-β R1/2) complex model showing a single binding site, alongside the corresponding PAE matrix. **E-F** Predicted U8–protein (TGF-β R1/2) complex model showing a single binding site, alongside the corresponding PAE matrix.

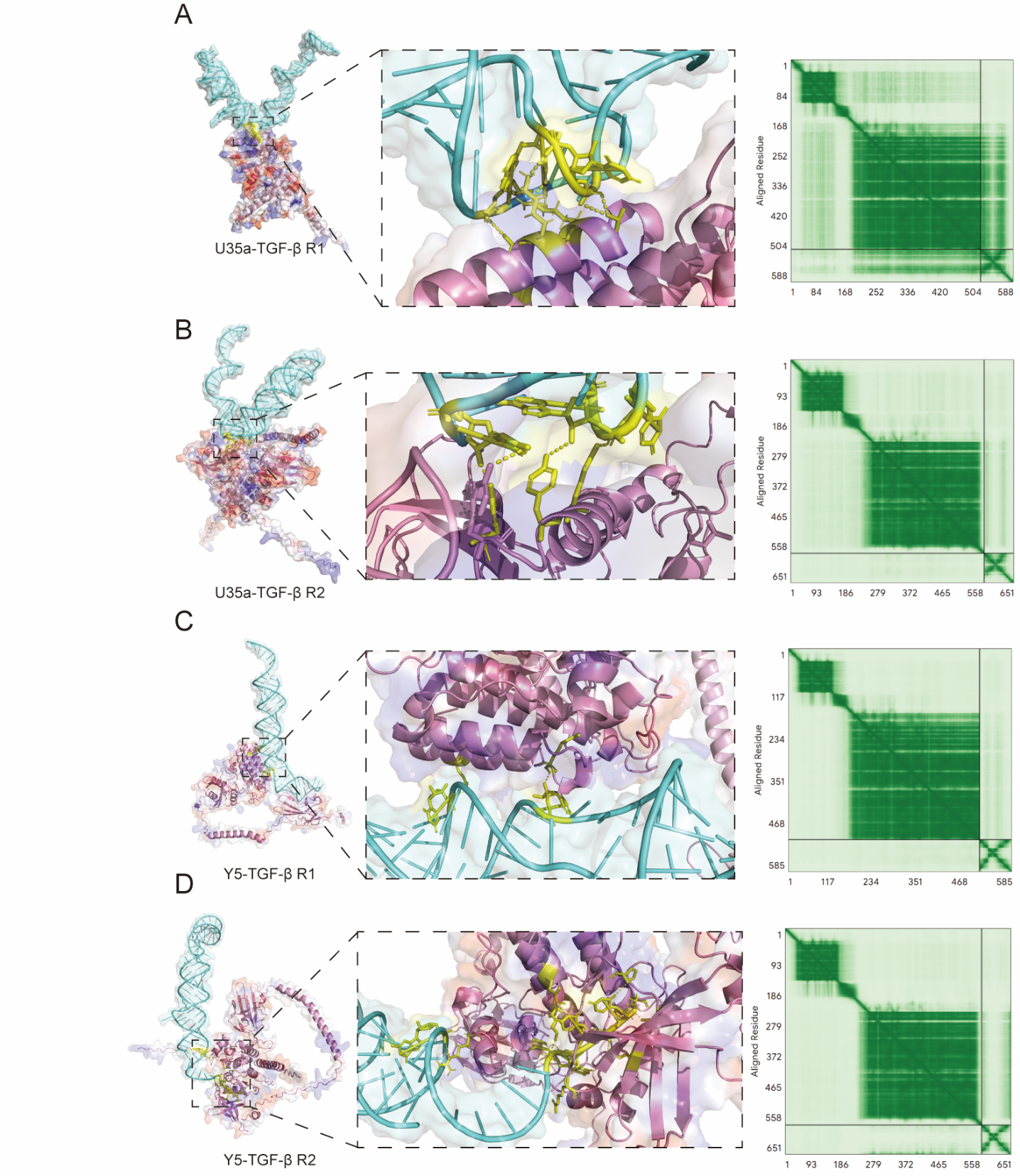

**Figure S2. AlphaFold 3 predictions of RNA–protein complexes.** Proteins and RNA are rendered as pink and cyan ribbons, respectively. Interacting amino acid residues and RNA bases are highlighted as yellow groups, with hydrogen bonds indicated by yellow dashed lines. On the electrostatic potential surface, blue and red patches represent densely positively and negatively charged regions, respectively. **A-B** Predicted U35a–protein (TGF-β R1/2) complex model showing a single binding site, alongside the corresponding predicted alignment error (PAE) matrix. **C-D** Predicted Y5–protein (TGF-β R1/2) complex model showing a single binding site, alongside the corresponding PAE matrix.

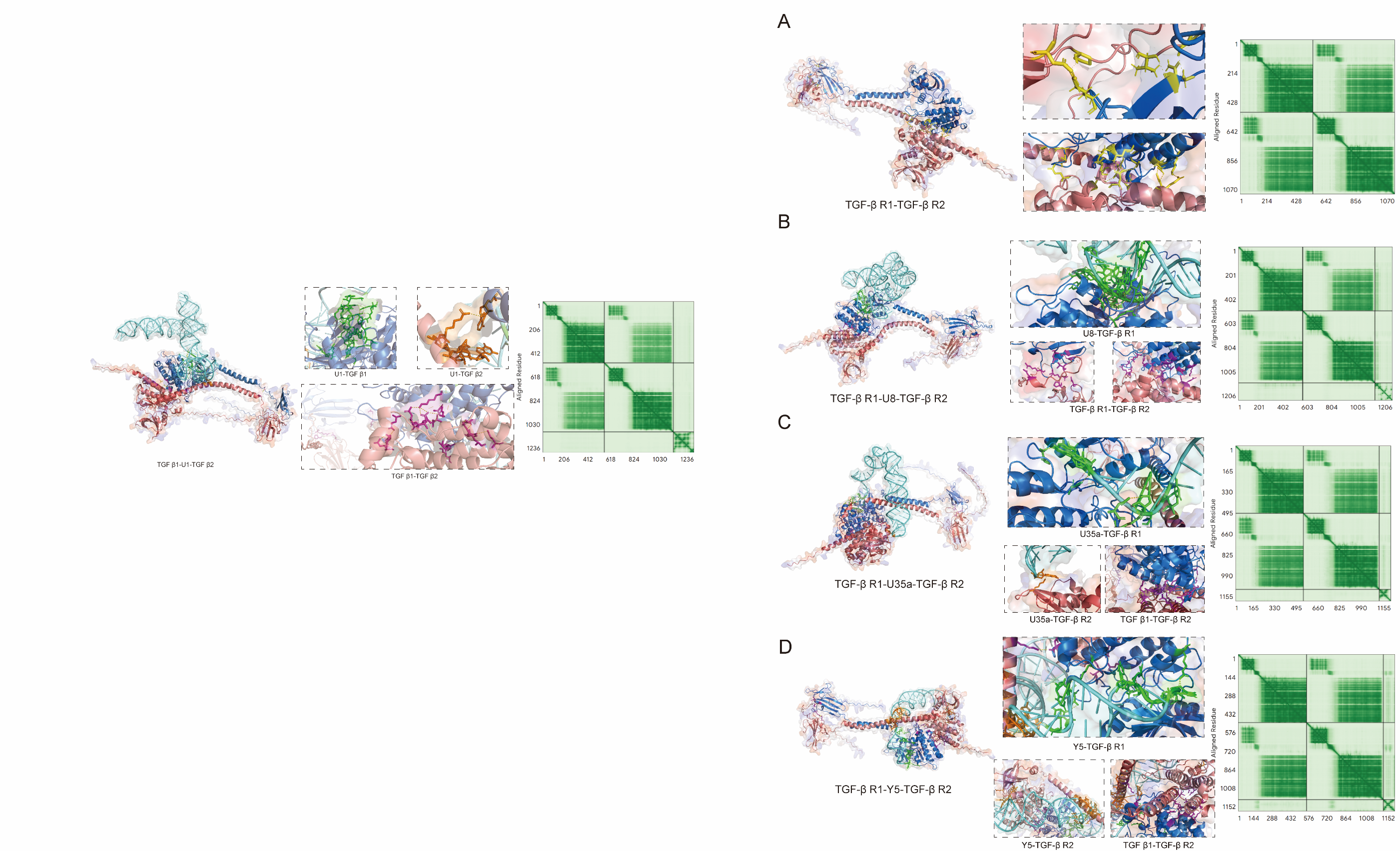

**Figure S3. AlphaFold 3 predictions of the TGF-β R1–RNA–TGF-β R2 ternary complex.** TGF-β R1 (blue), TGF-β R2 (red), and RNA (cyan) are shown in ribbon representation. Distinct interaction interfaces are color-coded as follows: RNA–TGF-β R1 (green), RNA–TGF-β R2 (orange), and TGF-β R1–TGF-β R2 (purple). Hydrogen bonds are denoted by yellow dashed lines. **A** Predicted TGF-β R1–TGF-β R2 interaction model, revealing two interfaces that link the homoproteins in head-to-head and tail-to-tail orientations, respectively, alongside the corresponding PAE matrix. **B–D** Predicted TGF-β R1–RNA–TGF-β R2 complex models displaying three types of interaction interfaces (RNA–TGF-β R1, RNA–TGF-β R2, and TGF-β R1–TGF-β R2) and the PAE matrix for the TGF-β R1–RNA–TGF-β R2 assembly.

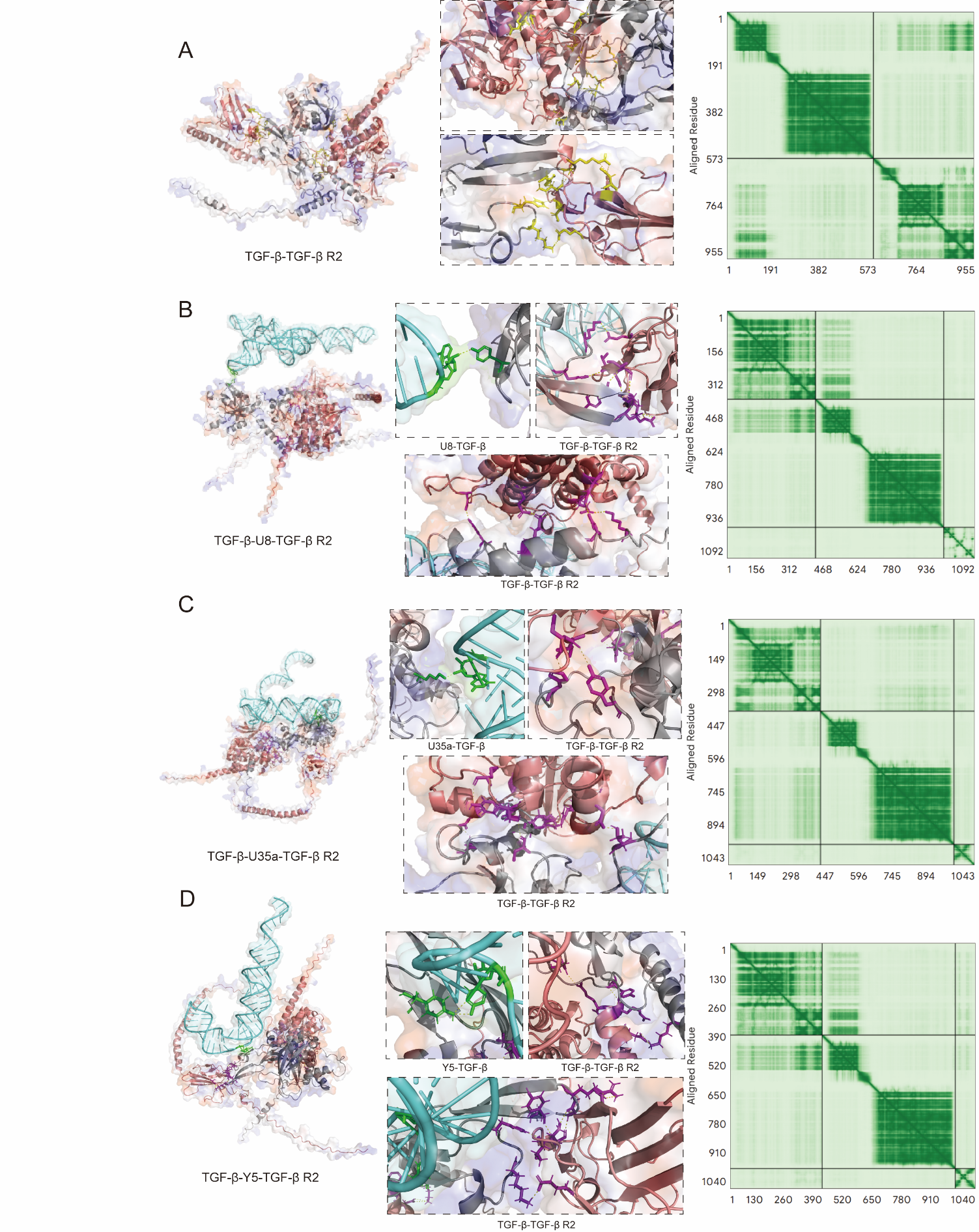

**Figure S4. AlphaFold 3 predictions of the TGF-β–RNA–TGF-β R2 ternary complex.** TGF-β (grey), TGF-β R2 (red), and RNA (cyan) are shown in ribbon representation. Distinct interaction interfaces in ternary complex are color-coded as follows: RNA–TGF-β (green) and TGF-β–TGF-β R2 (purple). Hydrogen bonds are denoted by yellow dashed lines. **A** Predicted TGF-β–TGF-β R2 interaction model, revealing that the two interfaces connect the two proteins in a disordered manner, alongside the corresponding PAE matrix. **B–D** Predicted TGF-β–RNA–TGF-β R2 complex models displaying two types of interaction interfaces (RNA–TGF-β and TGF-β R1–TGF-β R2) and the PAE matrix for the TGF-β–RNA–TGF-β R2 assembly.

**Table S1. Multicomponent complex interactions predicted by AlphaFold 3**

| Component | Ligand 1 | Ligand 2 | Hydrogen bond | ipTM | pTM |
| --- | --- | --- | --- | --- | --- |
| U1–TGF-β R1 | U1 | TGF-β R1 | 7 | 0.12 | 0.51 |
| U1–TGF-β R2 | U1 | TGF-β R2 | 9 | 0.10 | 0.47 |
| U3–TGF-β R1 | U3 | TGF-β R1 | 21 | 0.09 | 0.44 |
| U3–TGF-β R2 | U3 | TGF-β R2 | 0 | 0.17 | 0.47 |
| U8–TGF-β R1 | U8 | TGF-β R1 | 7 | 0.26 | 0.53 |
| U8–TGF-β R2 | U8 | TGF-β R2 | 8 | 0.28 | 0.51 |
| U35a–TGF-β R1 | U35a | TGF-β R1 | 5 | 0.42 | 0.63 |
| U35a–TGF-β R2 | U35a | TGF-β R2 | 4 | 0.12 | 0.52 |
| Y5–TGF-β R1 | Y5 | TGF-β R1 | 2 | 0.10 | 0.58 |
| Y5–TGF-β R2 | Y5 | TGF-β R2 | 9 | 0.25 | 0.53 |
| TGF-β R1-TGF-β R2 | TGF-β R1 | TGF-β R2 | 19 | 0.59 | 0.59 |
| TGF-β R1–U8–TGF-β R2 | U8 | TGF-β R1–TGF-β R2 | 33 | 0.50 | 0.54 |
|  | U8 | TGF-β R1 | 16 |  |  |
|  | U8 | TGF-β R2 | 0 |  |  |
|  | TGF-β R1 | TGF-β R2 | 17 |  |  |
| TGF-β R1–U35a–TGF-β R2 | U35a | TGF-β R1–TGF-β R2 | 25 | 0.52 | 0.55 |
|  | U35a | TGF-β R1 | 6 |  |  |
|  | U35a | TGF-β R2 | 1 |  |  |
|  | TGF-β R1 | TGF-β R2 | 18 |  |  |
| TGF-β R1–Y5–TGF-β R2 | Y5 | TGF-β R1–TGF-β R2 | 42 | 0.55 | 0.57 |
|  | Y5 | TGF-β R1 | 8 |  |  |
|  | Y5 | TGF-β R2 | 19 |  |  |
|  | TGF-β R1 | TGF-β R2 | 15 |  |  |

**Table S2. TGF-β–RNA–TGF-β R2 ternary complex interactions predicted by AlphaFold 3**

| Component | Ligand 1 | Ligand 2 | Hydrogen bond | ipTM | pTM |
| --- | --- | --- | --- | --- | --- |
| TGF-β–TGF-β R2 | TGF-β | TGF-β R2 | 17 | 0.32 | 0.39 |
| TGF-β–TGF-β R2-U8 | U8 | TGF-β–TGF-β R2 | 12 | 0.32 | 0.36 |
|  | U8 | TGF-β | 1 |  |  |
|  | U8 | TGF-β R2 | 0 |  |  |
|  | TGF-β | TGF-β R2 | 11 |  |  |
| TGF-β–TGF-β R2-U35a | U35a | TGF-β–TGF-β R2 | 12 | 0.18 | 0.40 |
|  | U35a | TGF-β | 1 |  |  |
|  | U35a | TGF-β R2 | 0 |  |  |
|  | TGF-β | TGF-β R2 | 11 |  |  |
| TGF-β–TGF-β R2-Y5 | Y5 | TGF-β–TGF-β R2 | 13 | 0.35 | 0.37 |
|  | Y5 | TGF-β | 2 |  |  |
|  | Y5 | TGF-β R2 | 0 |  |  |
|  | TGF-β | TGF-β R2 | 11 |  |  |

**Table S3. Protein sequence and AlphaFold 3 prediction**

|  | Name | Sequence | Reference | pTM |
| --- | --- | --- | --- | --- |
| Protein | TGF–bate receptor 1 | MEAAVAAPRPRLLLLVLAAAAAAAAALLPGATALQCFCHLCTKDNFTCVTDGLCFVSVTETTDKVIHNSMCIAEIDLIPRDRPFVCAPSSKTGSVTTTYCCNQDHCNKIELPTTVKSSPGLGPVELAAVIAGPVCFVCISLMLMVYICHNRTVIHHRVPNEEDPSLDRPFISEGTTLKDLIYDMTTSGSGSGLPLLVQRTIARTIVLQESIGKGRFGEVWRGKWRGEEVAVKIFSSREERSWFREAEIYQTVMLRHENILGFIAADNKDNGTWTQLWLVSDYHEHGSLFDYLNRYTVTVEGMIKLALSTASGLAHLHMEIVGTQGKPAIAHRDLKSKNILVKKNGTCCIADLGLAVRHDSATDTIDIAPNHRVGTKRYMAPEVLDDSINMKHFESFKRADIYAMGLVFWEIARRCSIGGIHEDYQLPYYDLVPSDPSVEEMRKVVCEQKLRPNIPNRWQSCEALRVMAKIMRECWYANGAARLTALRIKKTLSQLSQQEGIKM | P36897  (From Uniprot) | 0.66 |
|  | TGF–bate receptor 2 | MGRGLLRGLWPLHIVLWTRIASTIPPHVQKSVNNDMIVTDNNGAVKFPQLCKFCDVRFSTCDNQKSCMSNCSITSICEKPQEVCVAVWRKNDENITLETVCHDPKLPYHDFILEDAASPKCIMKEKKKPGETFFMCSCSSDECNDNIIFSEEYNTSNPDLLLVIFQVTGISLLPPLGVAISVIIIFYCYRVNRQQKLSSTWETGKTRKLMEFSEHCAIILEDDRSDISSTCANNINHNTELLPIELDTLVGKGRFAEVYKAKLKQNTSEQFETVAVKIFPYEEYASWKTEKDIFSDINLKHENILQFLTAEERKTELGKQYWLITAFHAKGNLQEYLTRHVISWEDLRKLGSSLARGIAHLHSDHTPCGRPKMPIVHRDLKSSNILVKNDLTCCLCDFGLSLRLDPTLSVDDLANSGQVGTARYMAPEVLESRMNLENVESFKQTDVYSMALVLWEMTSRCNAVGEVKDYEPPFGSKVREHPCVESMKDNVLRDRGRPEIPSFWLNHQGIQMVCETLTECWDHDPEARLTAQCVAERFSELEHLDRLSGRSCSEEKIPEDGSLNTTK | P37173  (From Uniprot) | 0.59 |
|  | TGF–bate 1 | MPPSGLRLLPLLLPLLWLLVLTPGRPAAGLSTCKTIDMELVKRKRIEAIRGQILSKLRLASPPSQGEVPPGPLPEAVLALYNSTRDRVAGESAEPEPEPEADYYAKEVTRVLMVETHNEIYDKFKQSTHSIYMFFNTSELREAVPEPVLLSRAELRLLRLKLKVEQHVELYQKYSNNSWRYLSNRLLAPSDSPEWLSFDVTGVVRQWLSRGGEIEGFRLSAHCSCDSRDNTLQVDINGFTTGRRGDLATIHGMNRPFLLLMATPLERAQHLQSSRHRRALDTNYCFSSTEKNCCVRQLYIDFRKDLGWKWIHEPKGYHANFCLGPCPYIWSLDTQYSKVLALYNQHNPGASAAPCCVPQALEPLPIVYYVGRKPKVEQLSNMIVRSCKCS | P01137  (From Uniprot) | 0.61 |

**Table S4. RNA sequence and AlphaFold 3 prediction**

|  | Name | Sequence | Reference | pTM |
| --- | --- | --- | --- | --- |
| RNA | **U1:** Homo sapiens (human) RNA, U1 small nuclear 1 (RNU1–1) | UUUCAUACUUACCUGGCAGGGGAGAUACCAUGAUCACGAAGGUGGUUUUCCCAGGGCGAGGCUUAUCCAUUGCACUCCGGAUGUGCUGACCCCUGCGAUUUCCCCAAAUGUGGGAAACUCGACUGCAUAAUUUGUGGUAGUGGGGGACUGCGUUCGCGCUUUCCCCUG | NR_004430.3 | 0.32 |
|  | **U3:** Homo sapiens (human) small nucleolar RNA, C/D box 3A (SNORD3A) | AAGACUAUACUUUCAGGGAUCAUUUCUAUAGUGUGUUACUAGAGAAGUUUCUCUGAACGUGUAGAGCACCGAAAACCACGAGGAAGAGAGGUAGCGUUUUCUCCUGAGCGUGAAGCCGGCUUUCUGGCGUUGCUUGGCUGCAACUGCCGUCAGCCAUUGAUGAUCGUUCUUCUCUCCGUAUUGGGGAGUGAGAGGGAGAGAACGCGGUCUGAGUGGU | NR_006880.1 | 0.25 |
|  | **U8:** Homo sapiens (human) U8 small nucleolar RNA | UUCUCAUGUGGGAUAGUUUUCACUUGUUCCUUCCUUUGGAGGGCAGAUUAGAACAUGAUGAAUGUAGUUUGCACAAUACAUUAUCAACAUCUUGGAGUUGUCAGAACUUGCAAUGCCCUGAUUUCGUUCUAUUUAC | Rfam: RF00096 | 0.20 |
|  | **Y5:** Homo sapiens (human) RNA, Ro60–associated Y5 (RNY5) | AGUUGGUCCGAGUGUUGUGGGUUAUUGUUAAGUUGAUUUAACAUUGUCUCCCCCCACAACCGCGCUUGACUAGCUUGCUGUUUU | NR_001571.2 | 0.23 |
|  | **U35a:** Homo sapiens (human) small nucleolar RNA, C/D box 35A (SNORD35A) | GGCAGAUGAUGUCCUUAUCUCACGAUGGUCUGCGGAUGUCCCUGUGGGAAUGGCGACAAUGCCAAUGGCUUAGCUGAUGCCAGGAG | NR_000018.1 | 0.22 |
